# Longitudinal changes in diet cause repeatable and largely reversible shifts in gut microbial communities of laboratory mice and are observed across segments of the entire intestinal tract

**DOI:** 10.1101/2021.05.06.443038

**Authors:** Adrian Low, Melissa Soh, Sou Miyake, Vanessa Zhi Jie Aw, Jian Feng, Adeline Wong, Henning Seedorf

## Abstract

Dietary changes are known to alter the composition of the gut microbiome. However, it is less understood how repeatable and reversible these changes are and how diet-switches affect the microbiota in the various segments of the gastrointestinal tract. Here, a treatment group of conventionally raised laboratory mice is subjected to two periods of western diet (WD) interrupted by a period of standard diet (SD) of the same duration. Beta-diversity analyses show that diet-induced microbiota changes are largely reversible (*q* = 0.1501; PERMANOVA, weighted-UniFrac comparison of the treatment-SD group to the control-SD group) and repeatable (*q* = 0.032; PERMANOVA, weighted-UniFrac comparison of both WD treatments). Furthermore, we report that diet switches alter the gut microbiota composition along the length of the intestinal tract in a segment-specific manner, leading to gut segment-specific Firmicutes/Bacteroidota ratios. We identified prevalent and distinct Amplicon Sequencing Variants (ASVs), particularly in genera of the recently described *Muribaculaceae*, along the gut as well as ASVs that are differentially abundant between segments of treatment and control groups. Overall, this study provides insights into the reversibility of diet-induced microbiota changes and highlights the importance to expand sampling efforts beyond the collections of fecal samples to characterize diet-dependent and segment-specific microbiome differences.

## 1. Introduction

Mice are popular model organisms for gut microbiome research partially due to the anatomical similarities between their gastrointestinal tracts (GIT) and that of humans [1]. Both, conventionally raised mice and germ-free mice, have enabled unprecedented insights into host-microbe interactions, including demonstrated causality between microorganisms and specific host phenotypes and/or metabolic markers [2,3]. However, while conventionally raised mice have been extensively studied, most studies only examined the fecal microbiota, which may not be representative of the entire GIT. Studies have shown a high similarity between fecal and colonic microbiotas [4,5] and a greater dissimilarity between microbiota of the lower intestinal and upper intestinal tract [5]. Given the disparate functions between segments of the GIT [6], microbes may adapt differently to changing conditions in the host. Indeed, segment specific bacterial phylotypes have been identified in healthy C57BL/6 mice [5], but it is not well established how dietary changes, such as repetitive exposure to high sucrose/high fat substrates typical of western diet (WD), may affect the microbiota in the different segments of the GIT. This is specifically true for some of the dominant taxa, which are not well described yet or for which isolates have only recently been obtained, e.g. members of the *Muribaculaceae* (formerly also known as S24-7 family). This family has a high prevalence in the intestinal tract of rodents, where they often constitute the major component of the gut Bacteroidota [7]. Whilst their exact function in the gut is still unclear, determining the spatial distribution along the gut would provide insight into their adaptability within the murine gut. Hence, where possible, it appears to be warranted and needed to examine the microbiota of the entire GIT to improve our understanding of the complex, gut-segment-dependent, associations and interactions between microbes and their hosts.

Diet-switches have been shown to strongly and rapidly affect the composition of the fecal microbiome [8]. However, reversibility and repeatability of diet induced changes in the gut microbiota remain to be elucidated in greater detail. Previous studies suggested that the duration of the dietary treatment, and potentially the type of experimental diet, impact on the reversibility of the induced change [9-11]. Diets such as WD, can drastically alter the gut microbiota/microbiome into configurations often associated with undesirable phenotypes such as adiposity [12], increased susceptibility to diseases [10], reduced gut length and mass [13,14]. Reversing WD-induced changes to the gut microbiota composition via a second dietary switch using a low-fat plant-based diet have been shown to ameliorate some of those undesirable phenotypes [10,15,16]. However, it is often not apparent if a reversal after a second diet switch is complete and/or if a third diet change, using the same diet as in the first diet switch, would alter the microbiota into the same composition as after the first diet switch.

In this study, C57BL/6J mice were fed WD for two periods (28 days each), interceded by a standard diet (SD) period of the same length (Figure 1A). A highly resolving 16S rRNA gene amplicon sequencing variant (ASV) approach [17] allowed us to analyze the reversibility and repeatability of diet-induced changes in the fecal and GIT segment microbiota of WD- and SD-fed mice. We show that microbiota of GIT segments are differentially affected by dietary changes and that changes in the fecal microbiota are repeatable and largely reversible.

**Figure 1.**
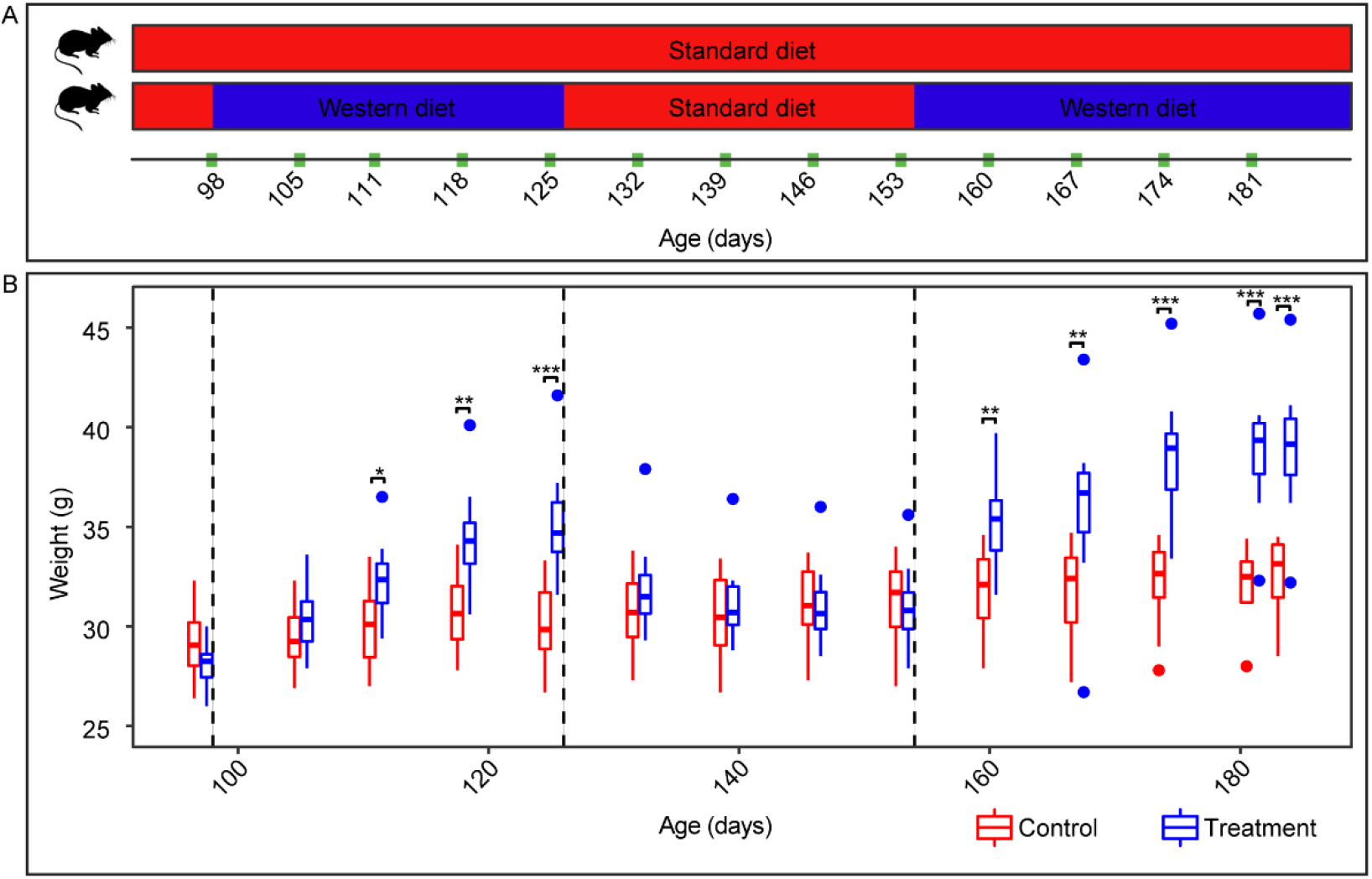
Experimental treatment and bodyweight over time. **(A)** Experimental timeline showing dietary regimens and timepoints for body weight measurements. **(B)** Boxplot depicts the weight of mice over time. Wilcoxon rank sum test was done to compare the weight between WD-and SD-mice (*n* = 12 mice per group). Asterisks represent significant difference where (*) denotes *q* < 0.05, (**) denotes *q* < 0.01 and (***) denotes *q* < 0.001. Dotted lines indicate a switch in diet for the treatment group.

## 2. Results

### 2.1 Effects of Western Diet on Animal Weight and Gastrointestinal Tract Length

All mice were weighed weekly to monitor bodyweight differences in response to the alternating diet regimen (Figure 1). Overall, the weight of the treatment mice (*n* = 12) exposed to two WD dietary regimens and one SD regimen was significantly different to that of the control group (*n* = 12) across 15 timepoints taken at weekly intervals (*q* = 4.63e−8, *N* = 360; Kruskal Wallis test; Figure 1B). Linear comparison showed that at the end of each WD regimen (day 27 and 83), the treatment mice weighed significantly more than SD-fed mice (*q*-values < 0.001; Wilcoxon rank sum test). Interestingly, the feeding of SD to treatment mice between the two WD periods led to a weight loss in the treatment group animals so that their bodyweight at the end of the SD period was indistinguishable to that of the animals in the control group. The control mice also gradually gained weight over 84 days.

To study the outcome of the dietary regimens on gut length, the postmortem length of GIT was compared between treatment and control groups (Table S1A for endpoint gut length and bodyweight). Comparison of the individual segment lengths revealed a significantly shorter GIT in the treatment group than in the control group (*q*-values < 0.05; Table S1B for Wilcoxon rank sum test). The same trend was observed after normalization of segment length to body weight (*q*-values < 0.05; Table S1C for Wilcoxon rank sum test).

### 2.2 Effects of Diet-Switches on the Fecal Microbiota

We compared weighted-UniFrac distances between treatment (WD-fed) and control groups (SD-fed), to analyze the effects of diet-switches on the fecal microbiota. We were particularly interested to determine how repeatable and reversible diet-induced changes of the fecal microbiome are. At the end of both WD regimens (day 27 and 83), there was -as expected-a significant separation between treatment and control groups (*q*-values < 0.001; Tables S2A for permutational multivariate analysis of variance (PERMANOVA) test; Figure 2A). However, the fecal microbiome configurations in the treatment group on days 27 and 83 were so similar that no significant difference was detectable (*q* = 0.0905; PERMANOVA), indicating a high repeatability of the diet-induced change. In contrast, fecal microbiota between the same timepoints differed significantly in the control group (*q* = 0.032; PERMANOVA) (Table S2A). We also observed that the diet-induced change is highly reversible as weighted-UniFrac distances between treatment and control groups on day 55 did not significantly differ (*q* = 0.1501; PERMANOVA, Figure 2A). It is noteworthy that Bray-Curtis dissimilarity indicated a significant difference between treatment and control group on day 55, showing that some differences between groups may persist even after four weeks (*q* = 0.0135; PERMANOVA). An ASV-level analysis of the diet-induced changes in the fecal microbiome is shown in Figure 2B (see Table S3A for mean relative abundance and taxonomic assignment of ASVs).

**Figure 2.**
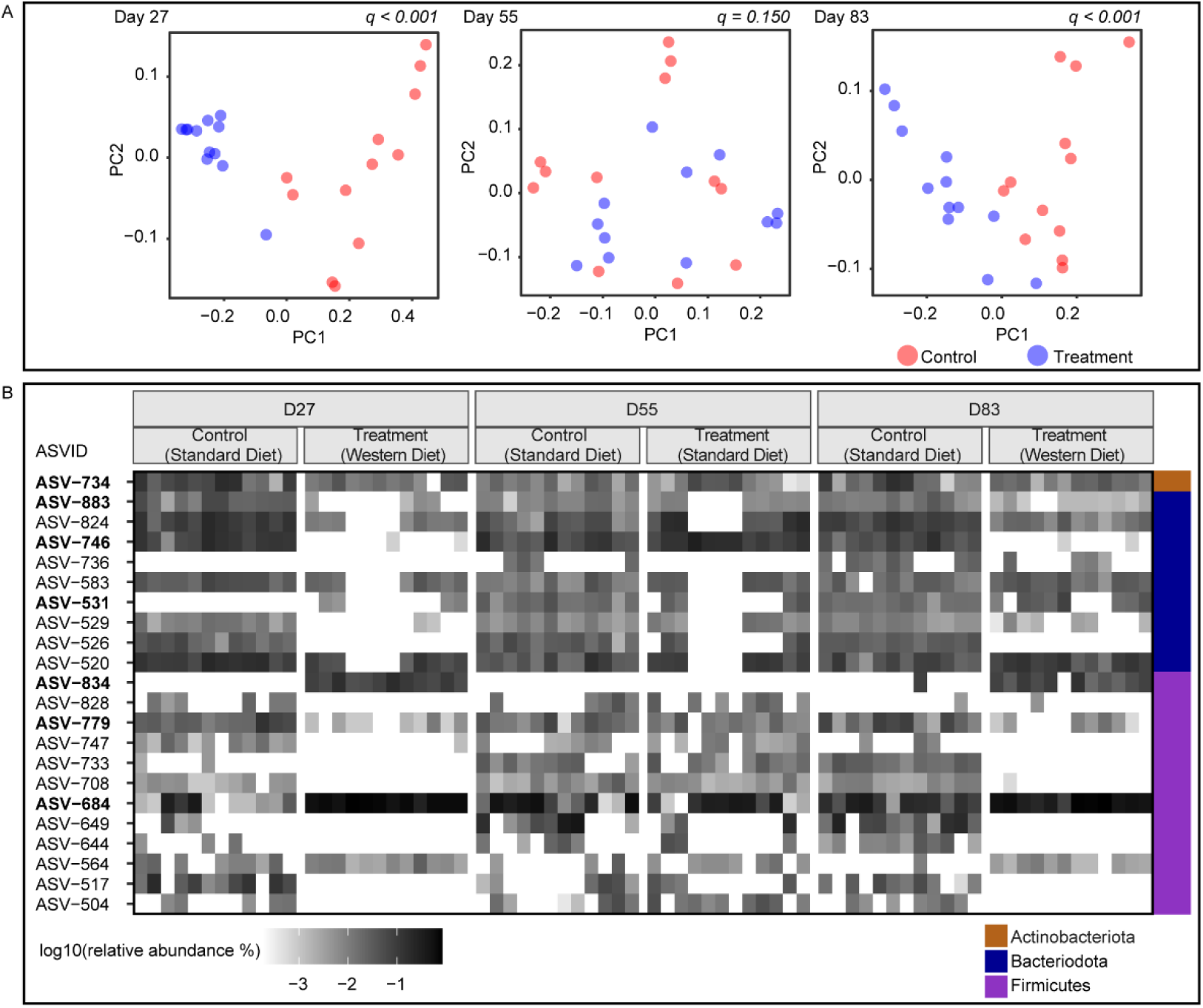
Beta-diversity between fecal microbiota of treatment (*n* = 12) and control (*n* = 12) groups. **(A)** Principal coordinate analysis (PCoA) plots for day-27, −55 and −83 based on the weighted-UniFrac distance. Statistical significance (*q*-values < 0.05) for PERMANOVA tests comparing weighted-UniFrac distance are shown. **(B)** A heatmap of 22 ASVs ≥ 0.5 relative abundance (5,203 counts per sample) per day in treatment and control. See Table S3A for taxonomic assignments. ASVs in bold have > 99% 16S rRNA gene similarity to the top BLASTn hit.

### 2.3 Diet-dependent Microbiota Differences are Observable along the Alimentary Tract

We compared alpha- and beta-diversity among the GIT segments (*N* = 140) and endpoint fecal samples (*N* = 24) to identify diet dependent and segmental differences between microbiotas. Overall, significantly reduced alpha-diversity was observed for most segments and fecal microbiotas of WD-fed mice compared to the control group according to Shannon (Figure 3A), Simpson’s (Figure 3B) and Chao1 (Figure 3C) indices. An exception to this was the ileal microbiota, where treatment and control groups shared similar species evenness (Shannon and Simpson’s indices). However, the ileum of WD-fed mice has significantly fewer rare (singletons and doubletons) ASVs than the control based on Chao1 index.

**Figure 3.**
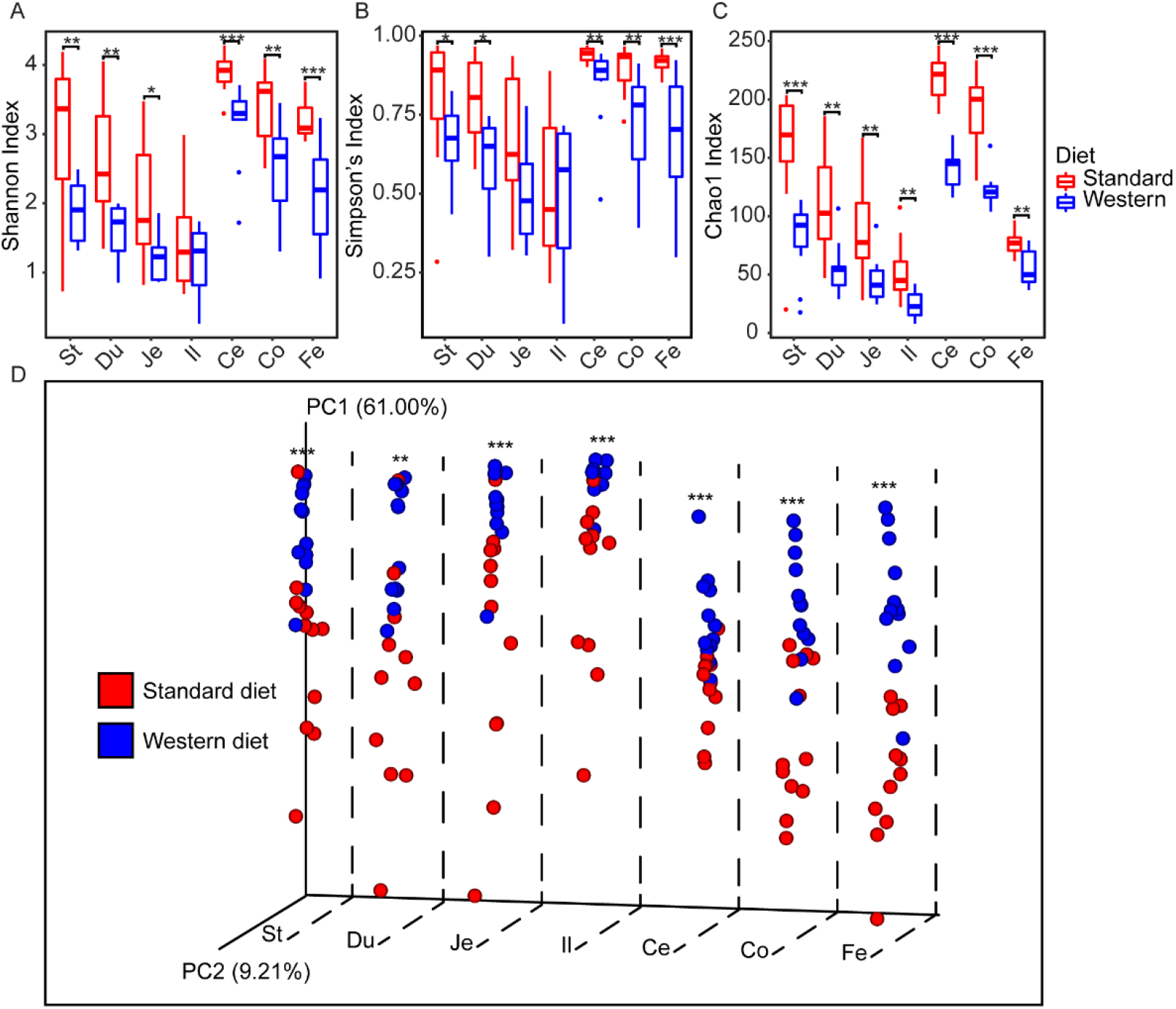
Alpha-and beta-diversity analyses of gut segment and fecal microbiota in western diet and standard diet-fed mice. Boxplots depict alpha-diversity shown as **(A)** Shannon, **(B)** Simpson and **(C)** Chao1 indices (Tables S2B-C for Wilcoxon rank sum test). **(D)** A principal coordinate analysis (PCoA) plot showing the weighted-UniFrac distance of grouped segments and fecal samples. Asterisks represent significant difference where (*) denotes *q* < 0.05, (**) denotes *q* < 0.01 and (***) denotes *q* < 0.001; PERMANOVA. St: stomach; Du: duodenum; Je: jejunum; Il: ileum; Ce: cecum; Co: colon; Fe: fecal.

Both Shannon and Simpson’s indexes decreased gradually from stomach to ileum with no statistically significant difference between adjacent segments (Figure 3A and 3B; *q*-values < 0.01; Tables S2B and S2C for Wilcoxon rank sum tests of Shannon and Simpson’s index, respectively). A significant increase was observed from ileum (least diverse) to the cecum (most diverse) followed by a decrease in the colon (*q*-values < 0.05; significant for Shannon index only). The Chao1 index followed a similar trend but with greater distinction between all segments of the treatment mice (Figure 3C; Table S2D for Wilcoxon rank sum test).

The comparison between fecal and segment microbiotas revealed different degrees of similarity in alpha diversity depending on the metric used. The treatment group fecal microbiota was highly similar to nearly all segments, except the cecum when using Simpson’s index (Table S2C). In contrast, fecal microbiota shared similar alpha diversity with fewer segments when using Shannon or Chao1 index (Shannon: stomach, duodenum and colon, Chao1: duodenum and jejunum; Tables S2B and S2D). The control fecal microbiota had similar alpha-diversity to the stomach (all three alpha-diversity indices), duodenum (all three alpha-diversity indices) and colon (Shannon and Simpson’s diversity).

Beta-diversity of gut segment microbiota was analyzed using Principal Coordinate Analysis (PCoA) of weighted-UniFrac distances, which showed statistically significant separation between treatment and control groups (Figure 3D; Table S4A for PERMANOVA). Longitudinal comparison within WD-fed mice showed that adjacent segments were significantly different to one another except jejunum vs ileum and cecum vs colon (Table S4B for PERMANOVA). In contrast, there was similarity between adjacent segments of the control group except ileum vs cecum and cecum vs colon (Table S4C for PERMANOVA). Fecal microbiotas were most similar to the respective colon microbiota in both groups (Tables S4B and S4C).

### 2.4 Taxonomic Compositional changes along the GIT and in Fecal Samples of WD-fed Mice

We analyzed microbiota at the phylum-, family- and ASV-level to determine compositional differences between WD-fed and SD-fed mice (Figure S1A, Figures 4 and 5, respectively). Firmicutes and Verrucomicrobiota (replaced Verrucomicrobia in SILVA 138) showed significantly higher mean relative abundances (Firmicutes = 86.7% ± 8.8 (Std. Dev.), *n* = 82; Verrucomicrobiota = 0.39% ± 0.74) in WD-fed mice than control mice (Firmicutes = 68.3% ± 17.5, *n* = 82; Verrucomicrobiota = 0.06% ± 0.23) (*q*-values < 0.001; Table S2E). Firmicutes were prevalent in all segments including feces in both groups. Verrucomicrobiota represented by *Akkermansiaceae* were more prevalent in the distal segments and feces compared to proximal segments of WD-fed mice. Other phyla showed significantly lower mean relative abundance than the control group (*q*-values < 0.001; Table S2E for Wilcoxon rank sum test). Exception to this is the phylum Proteobacteria, which did not differ significantly in mean relative abundances between the two groups (*q* = 0.31; Wilcoxon rank sum test). Consequently, the Firmicutes to Bacteroidota ratios (FBR) in the six gut segments as well as fecal samples revealed significantly greater variation in the treatment group than in the control group (*q* = 8.71e-05; Kruskal-Wallis test; Figure S1B).

**Figure 4.**
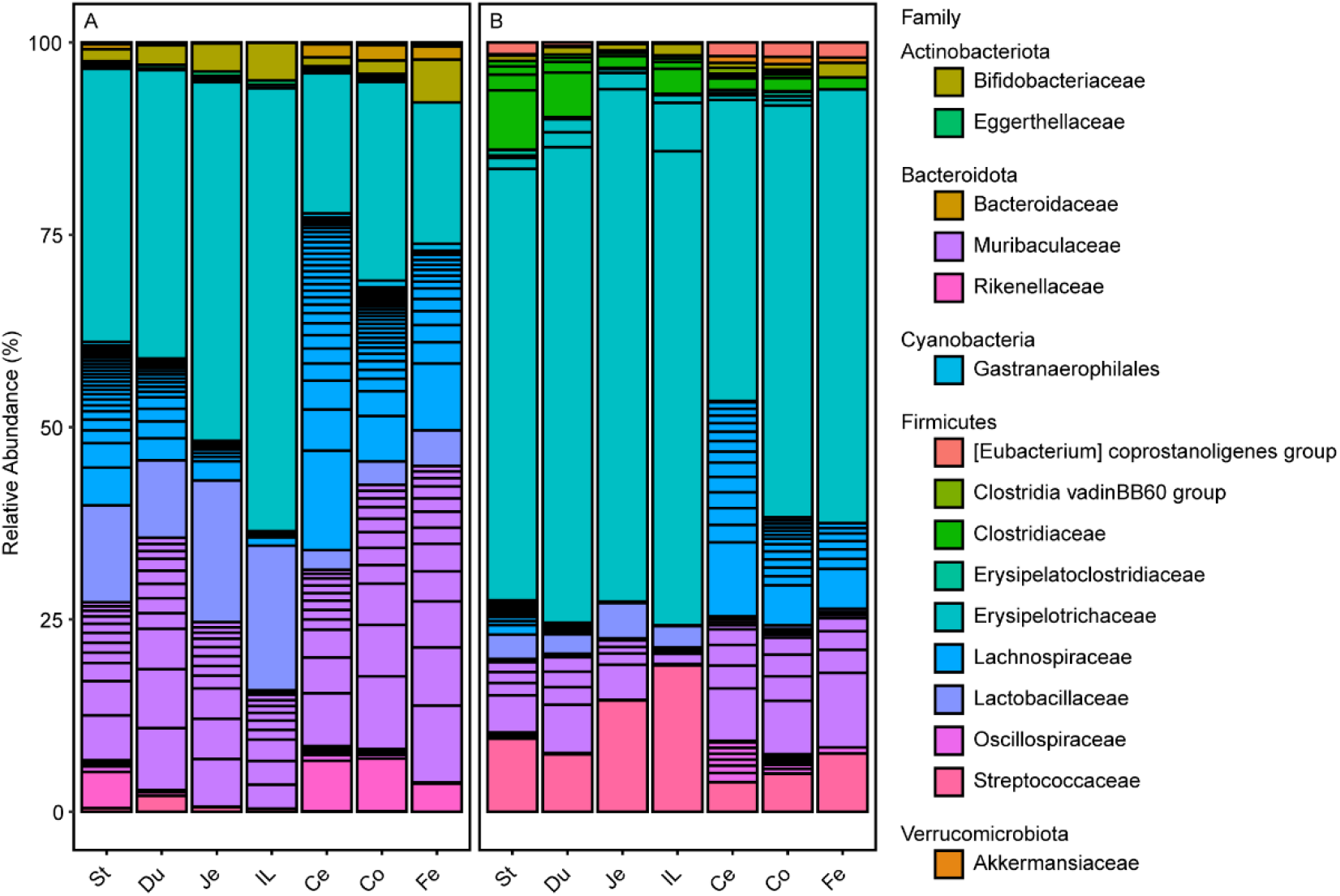
Gut microbiota classified at the family level. **(A)** Standard diet fed mice. **(B)** Western diet fed mice. Black horizontal lines on the bar-plots demarcate the different ASVs. Taxa are arranged alphabetically by phylum as indicated in legend headings. St: stomach; Du: duodenum; Je: jejunum; Il: ileum; Ce: cecum; Co: colon; Fe: fecal.

**Figure 5.**
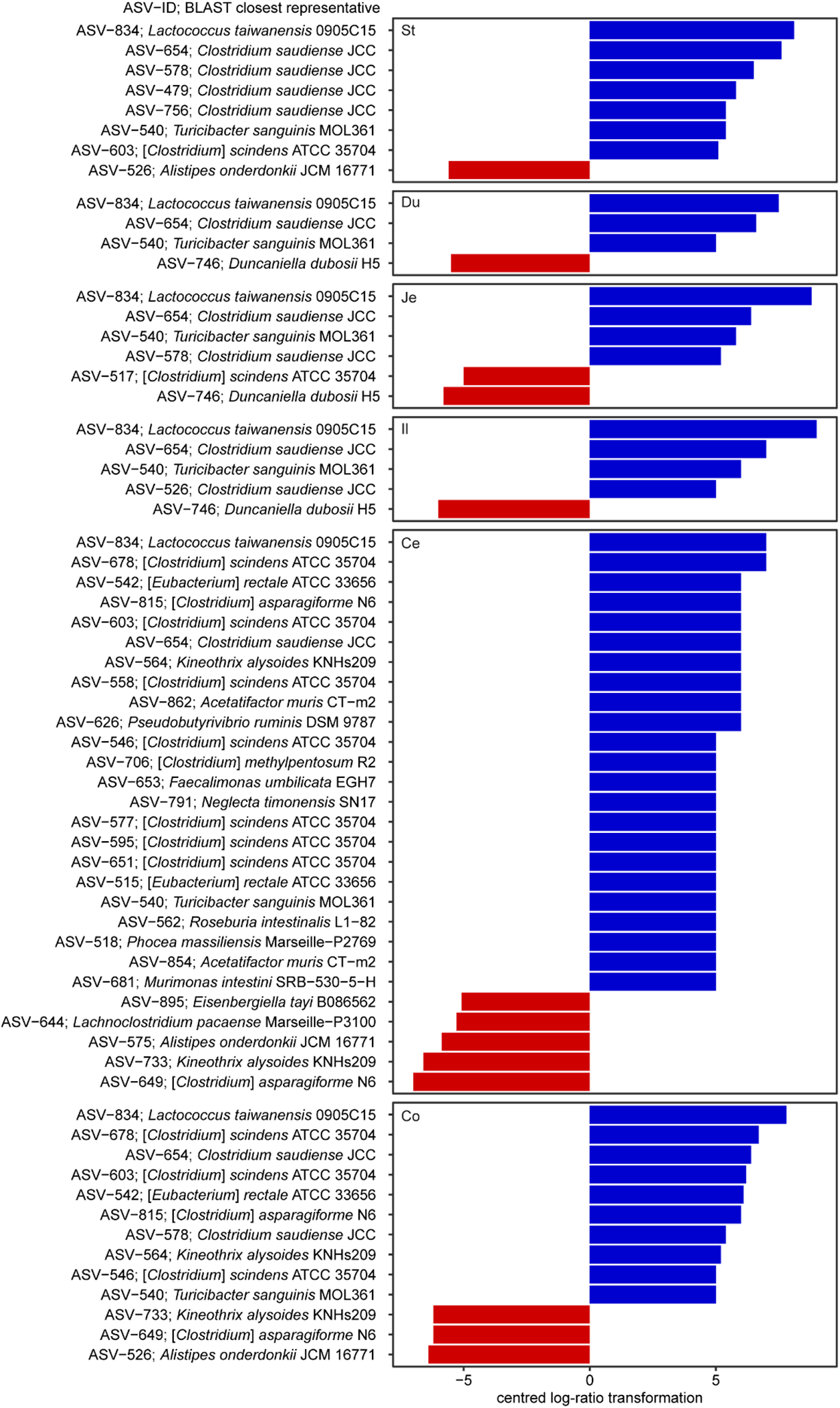
A bar-plot of differentially abundant ASVs within gut segments and fecal microbiota of mice fed standard or western diet. Standard diet and western diet are shown as red and blue, respectively. Differential abundant ASVs were identified using analysis of composition of microbiomes (ANCOM) method and have high *W* values are selected based on a distribution cutoff (Tables S5A for *W* values).

The high prevalence of the Firmicutes in WD-fed mice can be attributed to seven families (in order of highest to lowest relative abundance): *Erysipelotrichaceae, Streptococcaceae, Clostridiaceae*, [Eubacterium]_coprostanoligenes_group, *Enterococcaceae, Christensenellaceae* and *Staphylococcaceae*). These families have significantly higher mean relative abundances in WD-fed mice than in SD-fed mice (*q*-values < 0.001; Table S2F for Wilcoxon rank sum test and mean relative abundances). In contrast, the following nine families were significantly more abundant in SD-fed mice than in WD-fed mice (in order of highest to lowest relative abundance): *Lachnospiraceae, Lactobacillaceae*, Clostridia_vadinBB60_group, *Butyricicoccaceae, Acholeplasmataceae, Monoglobaceae*, RF39 and *Peptococcaceae* (*q*-values < 0.01; Table S2F). Five families were similarly abundant between the groups (in order of highest to lowest relative abundance): *Oscillospiraceae, Anaerovoracaceae, Ruminococcaceae, Erysipelatoclostridiaceae* and *Leuconostocaceae* (Table S2F).

ASV-level analysis revealed *Erysipelotrichaceae* ASV-684 as most abundant ASV in the treatment and control group. This ASV shares 100% identity to *Faecalibaculum rodentium* ALO17^T^. Moreover, analysis of composition of microbiomes (ANCOM) identified 69 ASVs that were differentially abundant in segments of treatment and control groups (Figure 5; Table S5A). Among these is a substantial fraction of ASVs that have low identity to cultured strains (median identity 94.2%). This includes ASVs of the family *Muribaculaceae. Muribaculaceae* have only been described recently and few isolates have been obtained thus far *[7,18-22]*. In this study we identified nine *Muribaculaceae* ASVs which have high abundance in either a gut segment or fecal sample of the treatment or control mice (Figure 6). Three of these ASVs (ASV-531 (*Paramuribaculum intestinale* B1404^T^), ASV-746 (*Duncaniella dubosii* H5^T^) and ASV-883 (*Muribaculum intestinale* YL27^T^)) shared identical 16S rRNA genes to known isolates while the remaining ASVs were low in similarity to cultured representatives (<96%, Table S3B). All nine ASVs were detectable along the GIT of control mice, following similar relative abundance patterns for the different segments. There were fewer ASVs of similar relative abundance in treatment mice than control mice (Figure 6). Notably, ASV-801 decreased to below detection in treatment mice. ASV-746, ASV-883 and ASV-736 decreased to <1% relative abundance throughout the GIT in treatment mice.

**Figure 6.**
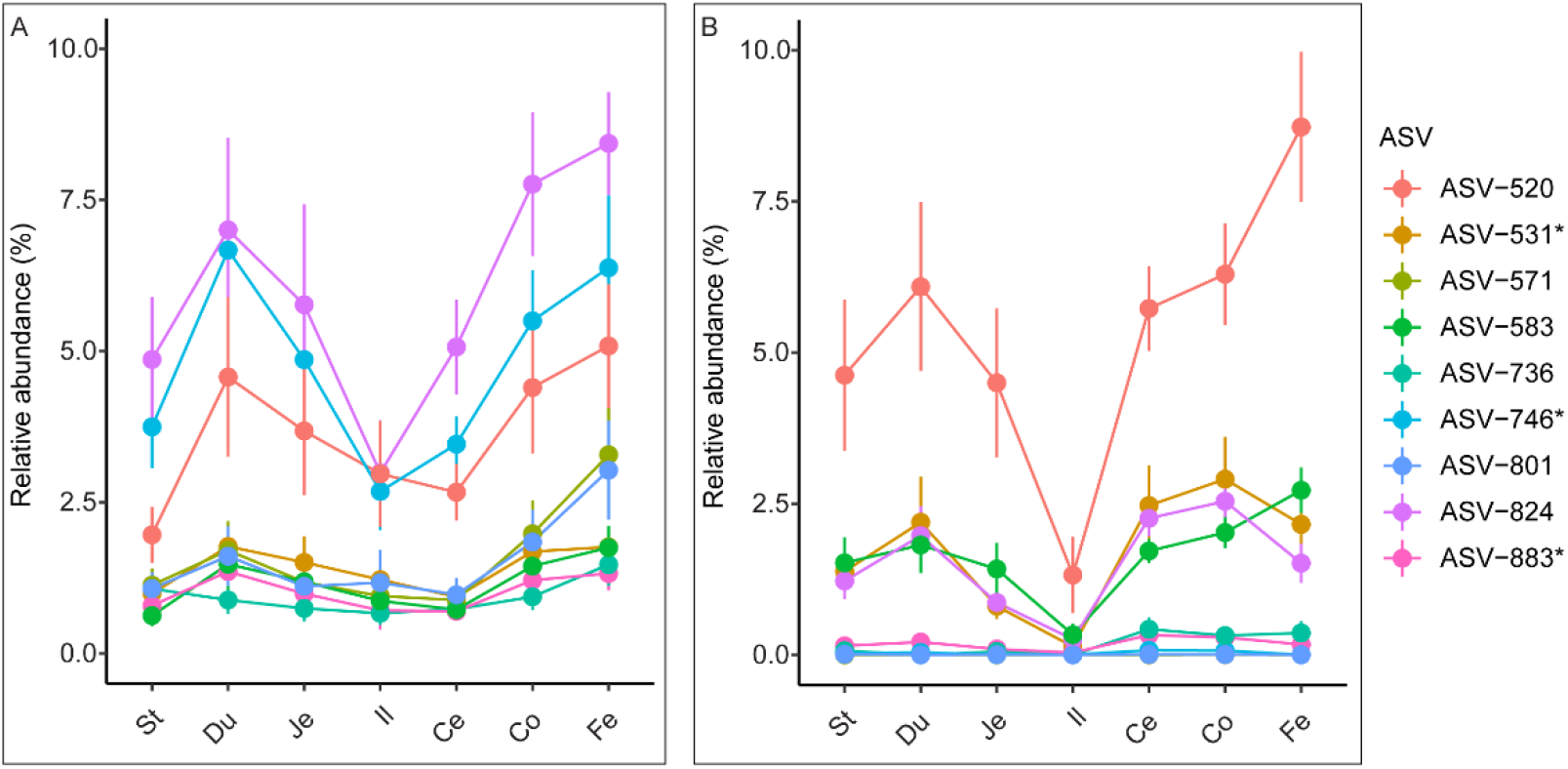
Relative abundance of nine *Muribaculaceae* ASVs along the gastrointestinal tract and feces. Shown are ASVs with ≥1% relative abundance in at least one segment/fecal group in **(A)** Control mice. **(B)** Treatment mice. Asterisks following ASVs indicate ASVs with identical 16S rRNA genes to type strains where ASV-531 is *Paramuribaculum intestinale* B1404^T^, ASV-746 is *Duncaniella dubosii* H5^T^ and ASV-883 is *Muribaculum intestinale* YL27^T^ (Table S3B). Error bars indicate standard error of the mean (*n* = 12 except duodenum: control *n* = 11; ileum: control *n* = 11, treatment *n* = 10). St: stomach; Du: duodenum; Je: jejunum; Il: ileum; Ce: cecum; Co: colon; Fe: fecal.

*Muribaculaceae* ASVs of control and treatment groups reached highest relative abundance in the duodenum of the upper intestinal tract and in the colon of the lower intestinal tract. In the treatment group, the lowest relative abundance of detectable *Muribaculaceae* ASVs was observed in the ileum, while control group *Muribaculaceae* ASVs were lowest in either ileum or cecum (Figure 6).

### 2.5 Predictive Metagenomic Profiling in Segments of Mice Fed Standard and Western Diets

The diet-induced changes in microbiota composition are also likely to affect the overall microbiome composition. PICRUSt2 was therefore used to predict E.C. enzymes and pathways from ASVs in gut segments. The weighted mean nearest sequenced taxon index (NSTI) scores were calculated as a measure of the mean phylogenetic distances of ASVs to their closest genomic representatives, i.e., accuracy of the prediction. The overall mean weighted NSTI was significantly lower for WD than SD fed mice (0.06 ± 0.04 (Std. Dev.); 0.10 ± 0.05, respectively; *n* = 70; *q* = 1e-05; Wilcoxon test). The NSTI scores for segments between treatment and control group were significantly different except within ileal microbiota (*q* = 0.11; Table S2H for Wilcoxon tests; Figure S3A). A Bray-Curtis dissimilarity-based non-metric multidimensional scaling (nMDS) of predicted enzyme counts revealed clustering by similar segment types with high similarity (90%) regardless of diet (Figure S3B). We observed enzymes of E.C. class 2 transferase and E.C. 3 hydrolase that highly correlated with segments of the small intestines (*rho* > 0.99; Spearman; *q*-values < 0.001; Table S4D for Wilcoxon rank sum test and mean relative abundance of E.C. between small and large intestines). Conversely, we observed E.C. class 1 oxidoreductase and E.C. class 4 lyase that highly correlated with cecal and colon samples (*rho* > 0.99; Spearman; *q*-values < 0.001; Table S4D).

Predicted MetaCyc pathways that are among the 20 most relatively abundant pathways in each segment revealed contrasting amino acid biosynthesis, metabolic and nucleotide salvage pathways between WD and SD groups (Figure S3C for heatmap). Notably, a sucrose degradation III pathway was significantly enriched in WD-fed mice (*q* = 3e-05; Wilcoxon rank sum test) from stomach to ileum with a mean relative abundance of 0.25% ± 0.06 (Std. Dev., *n* = 46) compared to the control (0.21% ± 0.04 (Std. Dev., *n* = 46)) (Figure S3C). In contrast, metabolic pathways of pyruvate fermentation to acetate and lactate II (*q* = 0.009; Wilcoxon rank sum test), pyruvate fermentation to isobutanol (*q* = 0.0003; Wilcoxon rank sum test) and galactose degradation (*q* = 0.036; Wilcoxon rank sum test) were significantly lower in the treatment than control group. ANCOM analysis of predicted MetaCyc pathways between treatment and control group revealed 17 pathways unique across the segments (Figure 7; Table S5B for *W* values). ANCOM identified MetaCyc pathways that were common from stomach to ileum of treatment mice: isopropanol biosynthesis, cob(II)yrinate a,c-diamide biosynthesis I (early cobalt insertion) and urea cycle. Five pathways common between the distal segments of treatment group were isopropanol biosynthesis, urea cycle, cob(II)yrinate a,c-diamide biosynthesis I (early cobalt insertion), peptidoglycan biosynthesis V (beta-lactam resistance) and peptidoglycan biosynthesis II (staphylococci). In the distal segments of control mice, common pathways were myo-, chiro- and scyllo-inositol degradation, superpathway of menaquinol-8 biosynthesis II, succinate fermentation to butanoate, 1,4-dihydroxy-6-naphthoate biosynthesis II and L-glutamate degradation V (via hydroxyglutarate). The pathway unique to the cecum and colon of control mice were 4-aminobutanoate degradation V and aromatic biogenic amine degradation (bacteria), respectively.

**Figure 7.**
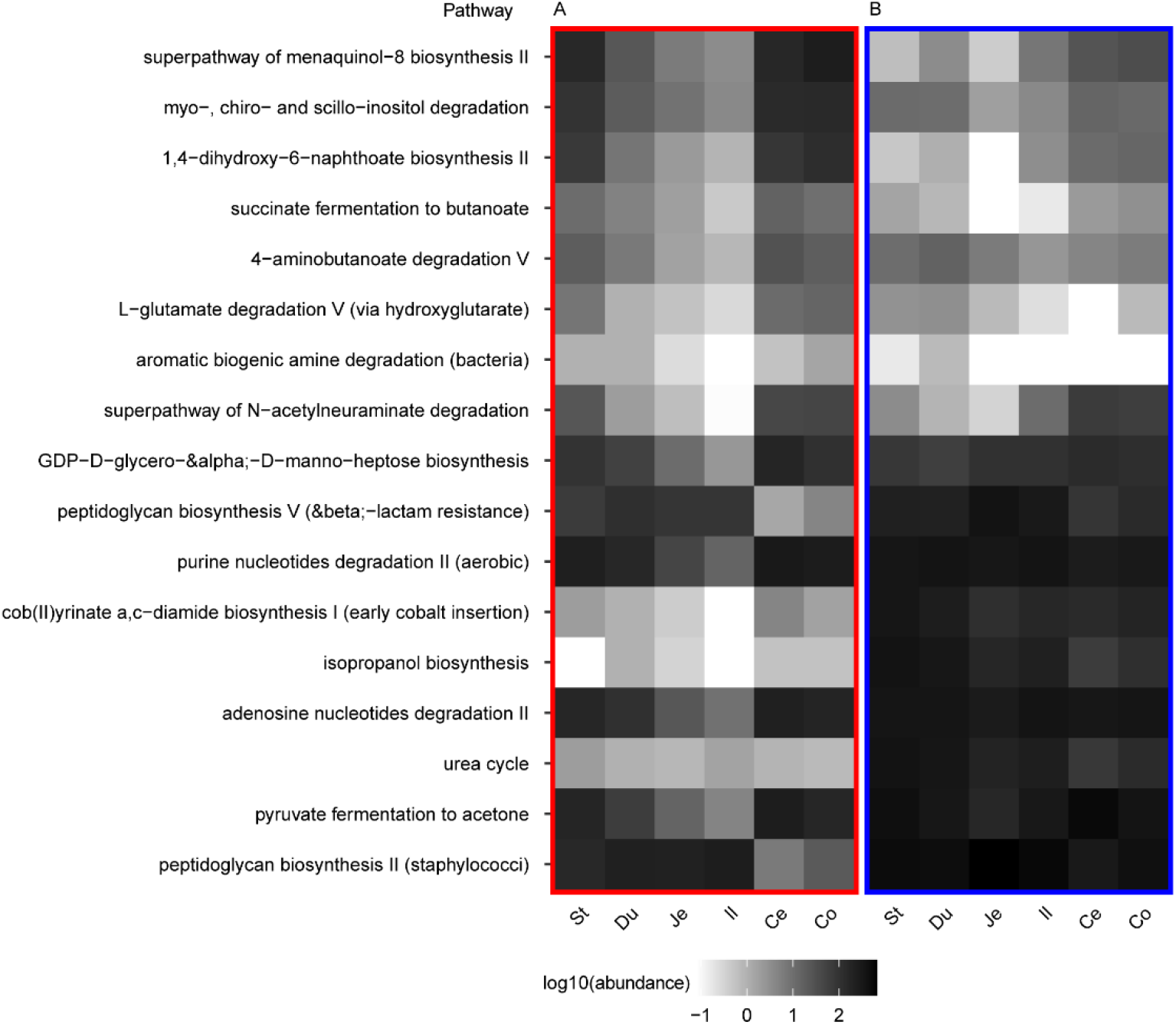
A heat-map representation of differentially abundant predicted pathways between segments of mice gastrointestinal tract of **(A)** standard or **(B)** western diet. Differential abundant MetaCyc pathways were identified using analysis of composition of microbiomes (ANCOM) method (Tables S5B for *W* values). The mean abundance (rarefied to 132,892 counts per sample) is shown. St: stomach; Du: duodenum; Je: jejunum; Il: ileum; Ce: cecum; Co: colon.

### 2.23. Discussion

In this study, we analyzed the gut microbiota of mice fed different dietary regimens to determine the effects of diet-switches on the animals, the reversibility and repeatability of diet-induced microbiota changes, and the GIT segment-specific microbiota compositions in response to a diet-switch experiment. The repeated feeding of WD had profound effects on the mice in that we observed shortened gut length and altered bodyweight. The changes to bodyweight and the GIT length of laboratory mice from dietary types have been previously reported to a similar extent [13]. The increase in bodyweight is primarily attributable to an increased intake of dietary calories, and to a lesser extent to low grade inflammation leading to insulin resistance [23]. However, factors causing a reduced GIT length from WD are less well understood. A lack of fermentable fiber in WD can reduce the mass and shorten cecal and colon lengths. These phenotypes may be ameliorated by short chain fatty acids (SCFAs) in the colon but not in cecal mass loss [24]. This indicates that different mechanisms for regulation of segment size and weight may exist, and that microorganisms could contribute to some of these mechanisms, e.g. via production of SCFAs as byproduct of fermentation of dietary fibers [25]. The mechanisms for the length phenotypes in other GIT segments remain to be elucidated in more detail.

We show that although dietary-switches induced largely reversible changes in the microbiota, a diet-switch may also lead to small-scale microbiota changes that persist even after extended periods of time, i.e. four weeks. This illustrates that diet not only rapidly changes the microbiota [8], but that dietary changes may also have long-lasting effects on the microbiome composition. From the obtained data it is not possible to infer if the microbiota would ultimately become entirely undistinguishable from the control mouse microbiota again or if the diet-switch induces changes that would persist permanently. A complete reversal of the gut microbiota may not be expected due to compositional variation among mice as they age and between cages [26,27]. Overall, our findings are consistent with previous studies indicating that WD induced microbiota changes to the gut microbiota may take a longer time to, or not fully reverse into the original state and may even be transmitted vertically to offspring [10]. It is also noteworthy that the diet composition may affect the impact and duration of the induced microbiota changes [9,14,16]. This is also evident in our study as we do not see significant microbiota differences between animals at the end of the two WD periods. The implications of these findings for diet-switch experiments in laboratory animals or for potential therapeutic interventions in humans warrant further investigations.

Lastly, we analyzed the effects of repeated feeding of WD on the composition of the microbiota and predicted microbiome in the different segments of the GIT. Regardless of the used alpha-diversity index used, alpha-diversity declined from stomach to ileum, then reaching a maximum in the cecum before decreasing again in the subsequent segments and feces. This is consistent with studies of laboratory and house mice, that observed that alpha-diversity is split anatomically between small and large intestines with significantly greater species richness and evenness in the latter segments, particularly the cecum [5,28,29]. These trends are independent of diet, potentially driven by multiple factors, such as coprophagy, transit time, etc.. However, it is also noteworthy that diet-specific effects impacted beta diversity. WD lowered beta diversity substantially so that even small microbial community structure differences between adjacent segments, e.g. duodenum and jejunum, were more noticeable than in SD-fed mice. Significant dissimilarities were also observed between fecal and stomach microbiota of WD-fed mice compared to the SD-fed control mice. This may indicate that the coprophagy behavior could be affected by the consumed diet. A reduced appetite for low fat/low sucrose diet has been observed in mice switching between WD and SD [11], which may suggest a dietary preference for WD and could provide an explanation for the observations. Ultimately, this change in eating behavior could also influence host physiology. Reduced coprophagy may not only affect the migration of fecal microbes into the gut but could limit nutrient intake such as vitamin B12 and folic acid that are beneficial to the host wellbeing [30,31].

An increased fecal Firmicutes/Bacteroidota ratio has been reported previously and appears to be a consistent response of the murine and human fecal microbiota to WD [32]. Our study reveals that the FBR varies between gut segments. Although higher variation and mean FBR between ileal microbiota and cecal or colon microbiota have been shown in C57BL/6J mice fed a refined high fat (60%) diet [14], similar comparisons to microbiotas of the upper intestinal tract have not been demonstrated. This is important as our study shows a greater shift towards a higher FBR in most of the microbiota of the upper intestinal tract compared to cecum and colon.

Of the Bacteroidota, the family *Muribaculaceae* was strongly impacted by WD and this study is -to our knowledge- the first to analyze the relative abundance of *Muribaculaceae* ASVs in the different segments of the intestinal tract. Diet differentially affected the relative abundance of the different *Muribaculaceae* ASVs, e.g. ASV-520 (currently uncultured) increased significantly in relative abundance in the treatment group. Currently, it remains difficult to deduce the ecological function of specific *Muribaculaceae* ASVs in this differential response. However, the results show that diet-induced microbiota changes could increase the relative abundance of uncultured microorganisms. This approach could be used to cultivate novel members of this family, either from feces or gut segments. A comparison of the spatial distribution of different *Muribaculaceae* species in the GITs of non-mouse hostsis currently difficult due to the lack of data on this recently described family. *Muribaculaceae* have been found predominantly in rodents but less so in humans or other animals, but it appears likely that each host type harbors specific *Muribaculaceae* species [7,28,33].

The exact role of Firmicutes ASVs in the murine intestinal tract remains -similarly to the beforementioned Bacteriodota-often not well understood. This is also the case for ASV-684 (100% identity to *Faecalibaculum rodentium*). ASV-684 is highly prevalent throughout the alimentary tract and feces during dietary changes. *Faecalibaculum rodentium* strains have been isolated from murine feces and the type strain has been shown to harbor genes associated with fermentation and alcohol utilization including L-lactate and butanol dehydrogenases [34,35]. To our knowledge, the extent of *F. rodentium* prevalence throughout the alimentary tract of laboratory murine gut has not been demonstrated before. However, it must be noted that experimental conditions, such as supplier of mice, may lead to observation of other phylotypes even when using C57BL/6 mice [5,36].

The use of PICRUSt2 to predict functional composition (E.C. numbers and MetaCyc pathways) revealed differences between proximal and distal segments. As monosaccharide absorption occurs mainly in the small intestine [37], correlation of enzymes involved in sugar or other substrate metabolism is consistent with the functional role of the small intestine. Similarly, as distal segments are more anoxic than proximal segments, enzymes involved in redox reactions such as dehydrogenases correlate with cecum and colon. Although a previous metagenomic study of fecal microbiomes revealed similar pathways and enzymes associated with simple sugar degradation in high fat/high sugar WD-fed mice [15], we demonstrate the spatial distribution of these enzymes within the gut. However, limitations to the accuracy of the presented predictions are given as the cultivation and characterizations of mouse microorganisms is -despite recent efforts-still not as comprehensive as databases available for human gut microbes. This is also shown by a study that compared the metagenomic profile of mice using a mouse reference database and PICRUSt2 default reference genomes, which indicated a better prediction using PICRUSt2 reference genomes [38,39]. Analytical tools for mouse gut microbiome research are likely to improve with the ongoing efforts to characterize the murine gut including metagenomic sequencing, isolation, and mechanistic studies [19,40].

In summary, our study shows that WD-induced microbiota changes are largely reversible and repeatable among the more abundant ASVs. Diet strongly alters relative abundances of ASVs and metabolic pathways along the lumen of the alimentary tract in a diet-dependent and in a segment-specific manner. Furthermore, our study also shows that a considerable fraction of mouse gut microorganisms remains uncultured. Cultivating these microorganisms will be a prerequisite for gnotobiotic mouse experiments that aim at improving our understanding of diet-dependent and gut biogeographic microbiome differences on host physiology and immune maturation.

## 4. Materials and Methods

### 4.1 Diet Experiment, Fecal Sampling and Gastrointestinal Segments Harvest

Twenty-four ten-week-old male C57BL/6J mice were randomly distributed into six cages of four mice each and maintained on SD for 34 days before the diet experiment commenced. Half the mice were fed ad libitum WD (carbohydrate = 49.9%, protein = 17.4%, fat = 20.0%; AIN-76A; TestDiet, St Louis, MO, USA) for 28 days, SD (carbohydrate = 53.4%, protein = 21.0%, fat = 5.0%; PicoLab Rodent Diet 20; LabDiet, St Louis, MO, USA) for another 28 days, and then WD for the last 30 days before sacrifice. The remaining mice acted as a control group and were fed SD throughout. Fresh fecal samples were collected weekly after which the mice were weighed (Figure 1B).

Mice were euthanized using carbon dioxide and subsequent cervical dislocation. Segments of the intestinal tract were sampled according to an established standard protocol [41]. Length of GIT segments (stomach, small intestine, cecum and colon) were measured directly after euthanasia and dissection of individual animals. Stomach, cecum, and colon segments were grossly further subdivided into two equal parts, while the jejunum was divided into three equal parts. Individual segment or parts and fresh fecal pellets were collected into sterile 2 mL screw cap tubes, flash frozen in liquid nitrogen, and stored at −80° C before DNA extraction.

### 4.2 DNA Extraction, Indexing and Amplicon Sequencing

Cells were lysed from sampled material using a zirconia (0.1 mm sized beads) bead-beating phenol-chloroform DNA extraction method as previously described [42]. DNA yield and purified PCR products were quantified using Quant-it Picogreen (Thermo Fisher Scientific, OR, USA). PCRs were carried out in triplicates (plus one no template control per sample) using primers 515F and 806R for the V4-V5 regions of the 16S rRNA gene as described in the Earth Microbiome Project protocol (Tables S6A and S6B for fecal and segment barcodes) [43-45]. Each PCR contained 25 µL of 1 × Taq PCR Mastermix (Qiagen GmbH, Hilden, Germany), final concentration of 0.2 µM of each primer, ≤ 20 ng of DNA and molecular water. PCRs were carried out in 96 well plates on a Bio-Rad thermal cycler using the following conditions: 94° C for 3 min, 35 cycles of 94° C for 45s, 50° C for 60s, 72° C for 90s followed by a final elongation of 72° C for 10 min. Equimolar concentrations of PCR products were purified and pooled using a 0.8 volume of AMPure XP beads (Beckman Coulter Genomics, Danvers, MA, USA) and eluted with molecular water. Amplicons were sequenced using the 250 bp paired-end sequencing chemistry on Illumina MiSeq platforms at Axil Scientific Pte Ltd and Genome Institute of Singapore, respectively.

### 4.3 Amplicon Sequences Processing, Microbiota and Predicted Metagenome Analyses

Demultiplexed fastq files for segment and fecal samples were provided by the sequencing vendor. Fastq files were processed using QIIME 2 2020.2 release with default options unless stated otherwise [46]. Primer sequences were removed from paired-end reads using the “qiime cutadapt trim-paired” command [47]. Paired-end reads were denoised, trimmed, clustered de-novo and chimera checked using the “qiime dada2 denoise-paired” command [48] with options: “--p-trunc-len-f 180” and “--p-trunc-len-r 137” to truncate forward and reverse reads, respectively. To minimize PCR artifacts, ASVs in less than four samples and fewer than 12 total sequences were filtered using the “qiime feature-table filter-features” command. Paired-end reads of segment samples with multiple parts namely, stomach, jejunum, cecum and colon were grouped by taking the mean ASV count using the “qiime feature-table group” command with the “--p-mode mean-ceiling” option (Table S6C for segment sample identities of merged sample parts). ASVs were assigned taxonomic identities using the “qiime feature-classifier classify-sklearn” command against the latest SILVA SSU for V4 region release 138 non-redundant 99% identity database, which has major changes to taxonomic nomenclature and phylogenetic lineages [49]. ASVs classified as unassigned, mitochondria and chloroplast were removed before further analysis. After quality filtering, the segment sample mean count was 33,641 reads per sample ± 10,519 (Std. Dev., *N* = 140) with 477 ASVs of 252 bp ± 0.44 (Std. Dev.) while fecal sample mean count was 9,935 reads per sample ± 3,889 (Std. Dev., *N* = 72) with 384 ASVs of 207 bp ± 0.40 (Std. Dev.) (Tables S6A and S6C for fecal and segment read count). We combined endpoint segment and fecal quality filtered feature tables and representative sequence files using “qiime feature-table merge” and “qiime feature-table merge-seqs”, respectively. The merged representative sequences were exported, aligned using the MUSCLE alignment tool and ends were trimmed to 201 bps using MEGA X version 10.2.0 [50,51]. The aligned fasta file was imported into QIIME 2 and aligned masking was performed using the “qiime alignment mask” command with the following options “--p-max-gap-frequency 0.2” to retain columns with no more than 20% gaps and “--p-min-conservation 0.8” to retain columns with at least 80% nucleotide. Unrooted and rooted phylogenetic trees for UniFrac distances were generated using “qiime phylogeny fasttree” and “qiime phylogeny midpoint-root” commands, respectively. A maximum likelihood tree was generated using the masked aligned sequences with “qiime phylogeny raxml-rapid-bootstrap” command with the options “--p-seed 477” and “--p-rapid-bootstrap-seed 898” to construct a reproducible tree, “--p-bootstrap-replicates 1000” for 1,000 bootstrapping replicates and “--p-substitution-model GTRCAT” for the GTR-CAT tree model [52]. The maximum likelihood tree was used to find identical sequences between fecal and segment dataset and where the sequences are identical, the ASV identity from fecal dataset was used for consistency. PICRUSt2 was used to predict enzymes (E.C. numbers) and MetaCyc metabolic pathways [53,54] of segment microbiota via the default pipeline using the “picrust2_pipeline.py” script. Briefly the PICRUSt2 script ran the following commands, “place_seqs.py” aligned ASVs to reference phylogeny, “hsp.py” obtained normalized 16S rRNA gene copies based on predicted genome to calculate NSTI values, E.C. abundances per genome. The command “pathway_pipeline.py” predicted the MetaCyc pathways from E.C. numbers, “add_descriptions.py” was used to generate enzymes and pathways output files.

ASVs featured in heatmaps and ANCOM analyses were further annotated by taking the top BLASTn hit against the NCBI 16S rRNA gene database [55]. Phylogenetic lineages of *Muribaculaceae* and *Lactobacilli* were manually curated to identify recently recognized bacterial species or formal changes to nomenclature [7,19-21,56]. The “qiime emperor plot” command was used to generate custom PCoA plots of dissimilarity matrices with categorical groups on the x-axis against the first principal coordinate (PC1) on the y-axis and PC2 on the z-axis. Emperor plots were visualized at https://view.qiime2.org/ [57]. All alpha-diversity metrices, conventional PCoA plots, heatmaps, bar graphs and boxplots were generated using R and relevant R packages including ggplot2, phyloseq, reshape2, tidyverse and qiime2R [58-63].

### 4.4 Statistical Analysis

R was used to perform statistical operations for mean, standard deviation, Kruskal-Wallis [64] and Wilcoxon rank sum tests with Benjamini-Hochberg false discovery rate (FDR) [65,66] based on counts from rarefied tables of segment and fecal (2,945 counts per sample) and fecal only (5,203 counts per sample). Pairwise PERMANOVA test was performed using the “qiime diversity beta-group-significance” command at 9,999 permutations [67]. An FDR-corrected *p*-value (*q* < 0.05) was considered as statistically significant. A Bray-Curtis similarity matrix of square-root transformed values of a rarefied table (716,085 counts per sample) of E.C. was visualized on a nMDS plot using PRIMER-E version 6.1.16 [68]. Other PRIMER-E functions that used the Bray-Curtis similarity matrix of E.C enzymes were Spearman’s rank correlation test between enzymes and sample groups and the Similarity Profile Analysis (SIMPROF) to find significant (*p* < 0.05) clusters of samples using 999 permutations. Differential abundance tests were performed on centered-log transformed counts of ASVs and predicted MetaCyc pathways using ANCOM to compare between treatment and control groups [69].

## Supplementary Materials

The following are available online at www.mdpi.com/xxx/s1, **Figure S1:** Phylum level comparison across segments and fecal microbiota of SD and WD-fed mice. **Figure S2:** A heat-map showing mean relative abundance of ASV ≥ 1% relative abundance per segment/fecal microbiota per diet. **Figure S3:** Predicted metagenomic profiles between segments of standard diet and western diet fed mice., **Table S1:** (A) Weight and length of mice at the end of 84 days of the experiment. (B) Wilcoxon rank sum test on segment length. (C) Wilcoxon rank sum test on segment lengths normalized to animal weight. **Table S2:** (A) PERMANOVA tests of the weighted-UniFrac distance of fecal samples. (B) Wilcoxon rank sum test comparing Shannon index between different GIT segments and diets. (C) Wilcoxon rank sum test comparing Simpson index between different GIT segments and diets. (D) Wilcoxon rank sum test comparing Chao1 index between different GIT segments and diets. (E) Pairwise phylum level Wilcoxon rank sum test between treatment and control groups. (F) Family level Wilcoxon rank sum test between treatment and control groups. (G) Wilcoxon rank sum test between ratios of Firmicutes:Bacteroidota ASV counts. (H) Wilcoxon rank sum test of NSTI values between segments and diets. **Table S3:** (A) ASV ≥0.5% (5,603 counts per sample) relative abundance of fecal samples. (B) ASVs ≥1% (2,945 counts per sample) relative abundance in segmental and endpoint fecal microbiota. **Table S4:** (A) Pairwise PERMANOVA test comparing weighted-UniFrac distances between segments of western diet and standard diet. (B) Pairwise PERMANOVA test comparing weighted-UniFrac distances between segments of western diet. (C) Pairwise PERMANOVA test comparing weighted-UniFrac distances between segments of standard diet. (D) Wilcoxon rank sum test of E.C. numbers between small and large intestines. **Table S5:** (A) ANCOM analysis of differentially abundant ASVs within segments of treatment and control group. (B) ANCOM analysis of differentially abundant MetaCyc pathways within segments of treatment and control group. **Table S6:** (A) Mapping file showing metadata and number of quality filtered reads used for fecal sample analysis. (B) Mapping file showing metadata used for segment analysis. (C) Quality filtered and merged (mean) samples of the same parts and mouse identity used in the study.

## Author Contributions

Conceptualization, S.M., A.W., and H.S.; methodology, S.M., A.W. and H.S.; software, A.L and M.S.; validation, A.L., M.S. and H.S.; formal analysis, A.L., M.S. and H.S.; investigation, A.L., S.M., A.W., M.S., F.J., and H.S.; resources, A.L., M.S. and H.S.; data curation, A.L. S.M. and M.S.; writing—original draft preparation, V.Z.J.A., A.L., M.S. and H.S..; writing—review and editing, A.L., M.S. and H.S.; visualization, M.S. and H.S.; supervision, H.S.; project administration, H.S.; funding acquisition, H.S.. All authors have read and agreed to the published version of the manuscript.

## Funding

This research was funded by Temasek Life Sciences Laboratory.

## Institutional Review Board Statement

All experiments involving mice used protocols approved by the Institutional Animal Care and Use Committee (IACUC) number: TLL-17-018 in accordance with National Advisory Committee for Laboratory Animal Research guidelines and were performed at Temasek Life Sciences Laboratory (TLL) with supervision by trained veterinarians.

## Data Availability Statement

FastQ files were uploaded to NCBI GenBank under BioProject ID PRJNA503296 SRA accession numbers SRX4970431-SRX4970677 for segments and SAMN13908703-SAMN13908726, SAMN13908823-SAMN13908846, SAMN13908943-SAMN13908966 for fecal samples.

## Acknowledgments

We thank Amit Anand and Muhammad Khairillah Bin Nanwi at TLL biocomputing for bioinformatics support.

## Conflicts of Interest

The authors declare no conflict of interest.

## Notes

### Competing Interest Statement

The authors have declared no competing interest.

## References

1. Hugenholtz, F.; de Vos, W.M. Mouse models for human intestinal microbiota research: a critical evaluation. Cell Mol Life Sci 2018, 75, 149–160, doi:10.1007/s00018-017-2693-8s.

2. Pawlowski, S.W.; Calabrese, G.; Kolling, G.L.; Platts-Mills, J.; Freire, R.; Alcantara-Warren, C.; Liu, B.; Sartor, R.B.; Guerrant, R.L. Murine model of Clostridium difficile infection with aged gnotobiotic C57BL/6 mice and a BI/NAP1 strain. J Infect Dis 2010, 202, 1708–1712, doi:10.1086/657086.

3. Bloom, S.M.; Bijanki, V.N.; Nava, G.M.; Sun, L.; Malvin, N.P.; Donermeyer, D.L.; Dunne, W.M., Jr.; Allen, P.M.; Stappenbeck, T.S. Commensal Bacteroides species induce colitis in host-genotype-specific fashion in a mouse model of inflammatory bowel disease. Cell Host Microbe 2011, 9, 390–403, doi:10.1016/j.chom.2011.04.009.

4. Turnbaugh, P.J.; Ridaura, V.K.; Faith, J.J.; Rey, F.E.; Knight, R.; Gordon, J.I. The effect of diet on the human gut microbiome: a metagenomic analysis in humanized gnotobiotic mice. Sci Transl Med 2009, 1, 6ra14–16ra14, doi:10.1126/scitranslmed.3000322.

5. Gu, S.; Chen, D.; Zhang, J.N.; Lv, X.; Wang, K.; Duan, L.P.; Nie, Y.; Wu, X.L. Bacterial community mapping of the mouse gastrointestinal tract. PLoS One 2013, 8, e74957, doi:10.1371/journal.pone.0074957.

6. Martinez-Guryn, K.; Leone, V.; Chang, E.B. Regional diversity of the gastrointestinal microbiome. Cell Host Microbe 2019, 26, 314–324, doi:10.1016/j.chom.2019.08.011.

7. Lagkouvardos, I.; Lesker, T.R.; Hitch, T.C.A.; Galvez, E.J.C.; Smit, N.; Neuhaus, K.; Wang, J.; Baines, J.F.; Abt, B.; Stecher, B.; et al. Sequence and cultivation study of Muribaculaceae reveals novel species, host preference, and functional potential of this yet undescribed family. Microbiome 2019, 7, 28, doi:10.1186/s40168-019-0637-2.

8. David, L.A.; Maurice, C.F.; Carmody, R.N.; Gootenberg, D.B.; Button, J.E.; Wolfe, B.E.; Ling, A.V.; Devlin, A.S.; Varma, Y.; Fischbach, M.A. Diet rapidly and reproducibly alters the human gut microbiome. Nature 2014, 505, 559–563.

9. Yang, B.; Ye, C.; Yan, B.; He, X.; Xing, K. Assessing the influence of dietary history on gut microbiota. Curr Microbiol 2019, 76, 237–247, doi:10.1007/s00284-018-1616-8.

10. Myles, I.A.; Fontecilla, N.M.; Janelsins, B.M.; Vithayathil, P.J.; Segre, J.A.; Datta, S.K. Parental dietary fat intake alters offspring microbiome and immunity. J Immunol 2013, 191, 3200–3209, doi:10.4049/jimmunol.1301057.

11. Carmody, R.N.; Gerber, G.K.; Luevano, J.M., Jr.; Gatti, D.M.; Somes, L.; Svenson, K.L.; Turnbaugh, P.J. Diet dominates host genotype in shaping the murine gut microbiota. Cell Host Microbe 2015, 17, 72–84, doi:10.1016/j.chom.2014.11.010.

12. Hu, S.; Wang, L.; Yang, D.; Li, L.; Togo, J.; Wu, Y.; Liu, Q.; Li, B.; Li, M.; Wang, G.; et al. Dietary fat, but not protein or carbohydrate, regulates energy intake and causes adiposity in mice. Cell Metab 2018, 28, 415–431 e414, doi:10.1016/j.cmet.2018.06.010.

13. Naya, D.E.; Karasov, W.H.; Bozinovic, F. Phenotypic plasticity in laboratory mice and rats: a meta-analysis of current ideas on gut size flexibility. Evol Ecol Res 2007, 9, 1363–1374.

14. Dalby, M.J.; Ross, A.W.; Walker, A.W.; Morgan, P.J. Dietary uncoupling of gut microbiota and energy harvesting from obesity and glucose tolerance in mice. Cell Rep 2017, 21, 1521–1533, doi:10.1016/j.celrep.2017.10.056.

15. Turnbaugh, P.J.; Backhed, F.; Fulton, L.; Gordon, J.I. Diet-induced obesity is linked to marked but reversible alterations in the mouse distal gut microbiome. Cell Host Microbe 2008, 3, 213–223, doi:10.1016/j.chom.2008.02.015.

16. Zhang, C.; Zhang, M.; Pang, X.; Zhao, Y.; Wang, L.; Zhao, L. Structural resilience of the gut microbiota in adult mice under high-fat dietary perturbations. ISME J 2012, 6, 1848–1857, doi:10.1038/ismej.2012.27.

17. Callahan, B.J.; McMurdie, P.J.; Holmes, S.P. Exact sequence variants should replace operational taxonomic units in marker-gene data analysis. ISME J 2017, 11, 2639–2643, doi:10.1038/ismej.2017.119.

18. Park, J.K.; Chang, D.H.; Rhee, M.S.; Jeong, H.; Song, J.; Ku, B.J.; Kim, S.B.; Lee, M.; Kim, B.C. Heminiphilus faecis gen. nov., sp. nov., a member of the family Muribaculaceae, isolated from mouse faeces and emended description of the genus Muribaculum. Antonie Van Leeuwenhoek 2021, 114, 275–286, doi:10.1007/s10482-021-01521-x.

19. Lagkouvardos, I.; Pukall, R.; Abt, B.; Foesel, B.U.; Meier-Kolthoff, J.P.; Kumar, N.; Bresciani, A.; Martínez, I.; Just, S.; Ziegler, C. The Mouse Intestinal Bacterial Collection (miBC) provides host-specific insight into cultured diversity and functional potential of the gut microbiota. Nat Microbiol 2016, 1, 16131, doi:10.1038/nmicrobiol.2016.131.

20. Miyake, S.; Ding, Y.; Soh, M.; Low, A.; Seedorf, H. Cultivation and description of Duncaniella dubosii sp. nov., Duncaniella freteri sp. nov. and emended description of the species Duncaniella muris. Int J Syst Evol Microbiol 2020, 70, 3105–3110, doi:10.1099/ijsem.0.004137.

21. Miyake, S.; Ding, Y.; Soh, M.; Low, A.; Seedorf, H. Muribaculum gordoncarteri sp. nov., an anaerobic bacterium from the faeces of C57BL/6J mice. Int J Syst Evol Microbiol 2020, doi:10.1099/ijsem.0.004338.

22. Miyake, S.; Ding, Y.; Soh, M.; Seedorf, H. Complete genome sequence of Duncaniella muris strain B8, isolated from the feces of C57/BL6 mice. Microbiol Resour Announc 2019, 8, doi:10.1128/MRA.00566-19.

23. Backhed, F.; Manchester, J.K.; Semenkovich, C.F.; Gordon, J.I. Mechanisms underlying the resistance to diet-induced obesity in germ-free mice. Proc Natl Acad Sci U S A 2007, 104, 979–984, doi:10.1073/pnas.0605374104.

24. Chassaing, B.; Miles-Brown, J.; Pellizzon, M.; Ulman, E.; Ricci, M.; Zhang, L.; Patterson, A.D.; Vijay-Kumar, M.; Gewirtz, A.T. Lack of soluble fiber drives diet-induced adiposity in mice. Am J Physiol Gastrointest Liver Physiol 2015, 309, G528–G541, doi:10.1152/ajpgi.00172.2015.

25. Nicholson, J.K.; Holmes, E.; Kinross, J.; Burcelin, R.; Gibson, G.; Jia, W.; Pettersson, S. Host-gut microbiota metabolic interactions. Science 2012, 336, 1262–1267, doi:10.1126/science.1223813.

26. Hoy, Y.E.; Bik, E.M.; Lawley, T.D.; Holmes, S.P.; Monack, D.M.; Theriot, J.A.; Relman, D.A. Variation in taxonomic composition of the fecal microbiota in an inbred mouse strain across individuals and time. PLoS One 2015, 10, e0142825, doi:10.1371/journal.pone.0142825.

27. Low, A.; Soh, M.; Miyake, S.; Seedorf, H. Host-age prediction from fecal microbiome composition in laboratory mice. bioRxiv 2020, doi:10.1101/2020.12.04.412734.

28. Suzuki, T.A.; Nachman, M.W. Spatial heterogeneity of gut microbial composition along the gastrointestinal tract in natural populations of house mice. PLoS One 2016, 11, e0163720, doi:10.1371/journal.pone.0163720.

29. Ericsson, A.C.; Gagliardi, J.; Bouhan, D.; Spollen, W.G.; Givan, S.A.; Franklin, C.L. The influence of caging, bedding, and diet on the composition of the microbiota in different regions of the mouse gut. Sci Rep 2018, 8, 4065, doi:10.1038/s41598-018-21986-7.

30. Ebino, K.Y.; Suwa, T.; Kuwabara, Y.; Saito, T.R.; Takahashi, K.W. [Lifelong coprophagy in male mice]. Jikken Dobutsu 1987, 36, 273–276, doi:10.1538/expanim1978.36.3_273.

31. Bogatyrev, S.R.; Rolando, J.C.; Ismagilov, R.F. Self-reinoculation with fecal flora changes microbiota density and composition leading to an altered bile-acid profile in the mouse small intestine. Microbiome 2020, 8, 19, doi:10.1186/s40168-020-0785-4.

32. Bisanz, J.E.; Upadhyay, V.; Turnbaugh, J.A.; Ly, K.; Turnbaugh, P.J. Meta-analysis reveals reproducible gut microbiome alterations in response to a high-fat diet. Cell Host Microbe 2019, 26, 265–272, doi:10.1016/j.chom.2019.06.013.

33. Seedorf, H.; Griffin, N.W.; Ridaura, V.K.; Reyes, A.; Cheng, J.; Rey, F.E.; Smith, M.I.; Simon, G.M.; Scheffrahn, R.H.; Woebken, D.; et al. Bacteria from diverse habitats colonize and compete in the mouse gut. Cell 2014, 159, 253–266, doi:10.1016/j.cell.2014.09.008.

34. Sharma, V.; Smolin, J.; Nayak, J.; Ayala, J.E.; Scott, D.A.; Peterson, S.N.; Freeze, H.H. Mannose alters gut microbiome, prevents diet-induced obesity, and improves host metabolism. Cell Rep 2018, 24, 3087–3098, doi:10.1016/j.celrep.2018.08.064.

35. Chang, D.-H.; Rhee, M.-S.; Ahn, S.; Bang, B.-H.; Oh, J.E.; Lee, H.K.; Kim, B.-C. Faecalibaculum rodentium gen. nov., sp. nov., isolated from the faeces of a laboratory mouse. Antonie Van Leeuwenhoek 2015, 108, 1309–1318, doi:10.1007/s10482-015-0583-3.

36. Ericsson, A.C.; Davis, J.W.; Spollen, W.; Bivens, N.; Givan, S.; Hagan, C.E.; McIntosh, M.; Franklin, C.L. Effects of vendor and genetic background on the composition of the fecal microbiota of inbred mice. PloS One 2015, 10, e0116704, doi:10.1371/journal.pone.0116704.

37. Drozdowski, L.A.; Thomson, A.B. Intestinal sugar transport. World J Gastroenterol 2006, 12, 1657–1670, doi:10.3748/wjg.v12.i11.1657.

38. Lesker, T.R.; Durairaj, A.C.; Galvez, E.J.C.; Lagkouvardos, I.; Baines, J.F.; Clavel, T.; Sczyrba, A.; McHardy, A.C.; Strowig, T. An integrated metagenome catalog reveals new insights into the murine gut microbiome. Cell Rep 2020, 30, 2909–2922 e2906, doi:10.1016/j.celrep.2020.02.036.

39. Jiang, P.; Green, S.J.; Chlipala, G.E.; Turek, F.W.; Vitaterna, M.H. Reproducible changes in the gut microbiome suggest a shift in microbial and host metabolism during spaceflight. Microbiome 2019, 7, 113, doi:10.1186/s40168-019-0724-4.

40. Liu, C.; Zhou, N.; Du, M.-X.; Sun, Y.-T.; Wang, K.; Wang, Y.-J.; Li, D.-H.; Yu, H.-Y.; Song, Y.; Bai, B.-B. The Mouse Gut Microbial Biobank expands the coverage of cultured bacteria. Nat. Commun. 2020, 11, 1–12.

41. Ruehl-Fehlert, C.; Kittel, B.; Morawietz, G.; Deslex, P.; Keenan, C.; Mahrt, C.R.; Nolte, T.; Robinson, M.; Stuart, B.P.; Deschl, U.; et al. Revised guides for organ sampling and trimming in rats and mice--part 1. Exp Toxicol Pathol 2003, 55, 91–106.

42. Rius, A.G.; Kittelmann, S.; Macdonald, K.A.; Waghorn, G.C.; Janssen, P.H.; Sikkema, E. Nitrogen metabolism and rumen microbial enumeration in lactating cows with divergent residual feed intake fed high-digestibility pasture. J Dairy Sci 2012, 95, 5024–5034, doi:10.3168/jds.2012-5392.

43. Parada, A.E.; Needham, D.M.; Fuhrman, J.A. Every base matters: assessing small subunit rRNA primers for marine microbiomes with mock communities, time series and global field samples. Environ Microbiol 2016, 18, 1403–1414, doi:10.1111/1462-2920.13023.

44. Apprill, A.; McNally, S.; Parsons, R.; Weber, L. Minor revision to V4 region SSU rRNA 806R gene primer greatly increases detection of SAR11 bacterioplankton. Aquat Microb Ecol 2015, 75, 129–137, doi:10.3354/ame01753.

45. Thompson, L.R.; Sanders, J.G.; McDonald, D.; Amir, A.; Ladau, J.; Locey, K.J.; Prill, R.J.; Tripathi, A.; Gibbons, S.M.; Ackermann, G.; et al. A communal catalogue reveals Earth’s multiscale microbial diversity. Nature 2017, 551, 457–463, doi:10.1038/nature24621.

46. Bolyen, E.; Rideout, J.R.; Dillon, M.R.; Bokulich, N.A.; Abnet, C.C.; Al-Ghalith, G.A.; Alexander, H.; Alm, E.J.; Arumugam, M.; Asnicar, F.; et al. Reproducible, interactive, scalable and extensible microbiome data science using QIIME 2. Nat Biotechnol 2019, 37, 1091, doi:10.1038/s41587-019-0252-6.

47. Martin, M. Cutadapt removes adapter sequences from high-throughput sequencing reads. EMBnet. journal 2011, 17, 10–12.

48. Callahan, B.J.; McMurdie, P.J.; Rosen, M.J.; Han, A.W.; Johnson, A.J.; Holmes, S.P. DADA2: High-resolution sample inference from Illumina amplicon data. Nat Methods 2016, 13, 581–583, doi:10.1038/nmeth.3869.

49. Quast, C.; Pruesse, E.; Yilmaz, P.; Gerken, J.; Schweer, T.; Yarza, P.; Peplies, J.; Glockner, F.O. The SILVA ribosomal RNA gene database project: improved data processing and web-based tools. Nucleic Acids Res 2013, 41, D590–D596, doi:10.1093/nar/gks1219.

50. Edgar, R.C. MUSCLE: multiple sequence alignment with high accuracy and high throughput. Nucleic Acids Res 2004, 32, 1792–1797, doi:10.1093/nar/gkh340.

51. Kumar, S.; Stecher, G.; Li, M.; Knyaz, C.; Tamura, K. MEGA X: molecular evolutionary genetics analysis across computing platforms. Mol Biol Evol 2018, 35, 1547–1549.

52. Stamatakis, A. RAxML version 8: a tool for phylogenetic analysis and post-analysis of large phylogenies. Bioinformatics 2014, 30, 1312–1313, doi:10.1093/bioinformatics/btu033.

53. Caspi, R.; Foerster, H.; Fulcher, C.A.; Kaipa, P.; Krummenacker, M.; Latendresse, M.; Paley, S.; Rhee, S.Y.; Shearer, A.G.; Tissier, C. The MetaCyc Database of metabolic pathways and enzymes and the BioCyc collection of Pathway/Genome Databases. Nucleic Acids Res 2007, 36, D623–D631, doi:10.1093/nar/gkr1014.

54. Douglas, G.M.; Maffei, V.J.; Zaneveld, J.R.; Yurgel, S.N.; Brown, J.R.; Taylor, C.M.; Huttenhower, C.; Langille, M.G.I. PICRUSt2 for prediction of metagenome functions. Nat Biotechnol 2020, doi:10.1038/s41587-020-0548-6.

55. Zhang, Z.; Schwartz, S.; Wagner, L.; Miller, W. A greedy algorithm for aligning DNA sequences. J Comput Biol 2000, 7, 203–214, doi:10.1089/10665270050081478.

56. Zheng, J.; Wittouck, S.; Salvetti, E.; Franz, C.M.; Harris, H.M.; Mattarelli, P.; O’Toole, P.W.; Pot, B.; Vandamme, P.; Walter, J. A taxonomic note on the genus Lactobacillus: Description of 23 novel genera, emended description of the genus Lactobacillus Beijerinck 1901, and union of Lactobacillaceae and Leuconostocaceae. Int J Syst Evol Microbiol 2020, 70, 2782–2858.

57. Vazquez-Baeza, Y.; Gonzalez, A.; Smarr, L.; McDonald, D.; Morton, J.T.; Navas-Molina, J.A.; Knight, R. Bringing the dynamic microbiome to life with animations. Cell Host Microbe 2017, 21, 7–10, doi:10.1016/j.chom.2016.12.009.

58. Wickham, H. Reshaping data with the reshape package. J Stat Softw 2007, 21, 1–20.

59. Wickham, H. ggplot2: elegant graphics for data analysis. 2016.

60. R Core Team R: A language and environment for statistical computing, R Foundation for Statistical Computing: Vienna, Austria, 2013.

61. McMurdie, P.J.; Holmes, S. phyloseq: an R package for reproducible interactive analysis and graphics of microbiome census data. PloS one 2013, 8, e61217, doi:10.1371/journal.pone.0061217.

62. Bisanz, J.E. qiime2R: Importing QIIME2 artifacts and associated data into R sessions, v0.99; 2018.

63. Wickham, H.; Averick, M.; Bryan, J.; Chang, W.; McGowan, L.; François, R.; Grolemund, G.; Hayes, A.; Henry, L.; Hester, J. Welcome to the Tidyverse. J Open Source Softw 2019, 4, 1686.

64. Kruskal, W.H.; Wallis, W.A. Use of ranks in one-criterion variance analysis. J Am Stat Assoc 1952, 47, 583–621.

65. Benjamini, Y.; Hochberg, Y. Controlling the false discovery rate: a practical and powerful approach to multiple testing. J R Stat Soc Series B Stat Methodol 1995, 57, 289–300.

66. Wilcoxon, F. Individual comparisons by ranking methods. Biometrics 1945, 1, 80–83.

67. Anderson, M.J. A new method for non-parametric multivariate analysis of variance. Austral Ecol 2001, 26, 32–46, doi:10.1111/j.1442-9993.2001.01070.pp.x.

68. Clarke, K.R. Non-parametric multivariate analyses of changes in community structure. Aust J Ecol 1993, 18, 117–143, doi:10.1111/j.1442-9993.1993.tb00438.x.

69. Mandal, S.; Van Treuren, W.; White, R.A.; Eggesbo, M.; Knight, R.; Peddada, S.D. Analysis of composition of microbiomes: a novel method for studying microbial composition. Microb Ecol Health Dis 2015, 26, 27663, doi:10.3402/mehd.v26.27663.

